# Intrinsic Disorder and Salt-dependent Conformational Changes of the N-terminal Region of TFIP11 Splicing Factor

**DOI:** 10.1101/2024.02.18.580885

**Authors:** Blinera Juniku, Julien Mignon, Rachel Carême, Alexia Genco, Anna Maria Obeid, Denis Mottet, Antonio Monari, Catherine Michaux

**Affiliations:** Laboratory of Physical Chemistry of Biomolecules, UCPTS, University of Namur, Rue de Bruxelles 61, B-5000 Namur, Belgium; Namur Research Institute for Life Sciences (NARILIS), University of Namur, Namur, Belgium; Namur Institute of Structured Matter (NISM), University of Namur, Namur, Belgium; GIGA-Molecular Biology of Diseases, Molecular Analysis of Gene Expression (MAGE) Laboratory, University of Liege, B34, Avenue de l’Hôpital, B-4000 Liège, Belgium; Université Paris Cité and CNRS, ITODYS, F-75006, Paris, France

**Keywords:** Tuftelin Interacting Protein 11, spliceosome protein, intrinsically disordered protein, polyampholyte, spectroscopy, molecular dynamics, protein assembly

## Abstract

Tuftelin Interacting Protein 11 (TFIP11) was recently identified as a critical human spliceosome assembly regulator, interacting with multiple spliceosome proteins and localises in several membrane-less organelles. However, there is a lack of structural information on TFIP11, limiting the rationalisation of its biological role. TFIP11 has been predicted as a highly disordered protein, and more specifically concerning its N-terminal (N-TER) region. Intrinsically disordered proteins (IDPs) lack a defined tertiary structure, existing as a dynamic conformational ensemble, favouring their role as hubs in protein-protein and protein-RNA interaction networks. Furthermore, IDPs are involved in liquid-liquid phase separation (LLPS), driving the formation of subnuclear compartments.

Combining disorder prediction, molecular dynamics, and spectroscopy methods, this contribution shows the first evidence TFIP11 N-TER may be described as a polyampholytic IDP, exhibiting a structural duality with the coexistence of ordered and disordered assemblies, depending on the ionic strength of the protein environment. Increasing the salt concentration enhances the protein conformational flexibility, presenting a fuzzier conformational landscape, a more globule-like shape, and an unstructured arrangement that could favour LLPS segregation and protein-RNA interaction. The regions mostly composed of charged and hydrophilic residues are the most impacted, including the G-Patch domain which is of crucial importance to TFIP11 function.

## 1. Introduction

Splicing of precursor messenger RNA (pre-mRNA) is a fundamental process in eukaryotic gene expression. The splicing of introns in pre-mRNA is carried out by the spliceosome, a dynamic macromolecular complex composed of five small nuclear RNA (snRNAs U1, U2, U4, U5, and U6) associated with more than 200 proteins. Due to its crucial role in RNA processing, spliceosome is a key cellular machinery in regulating gene expression, and its deregulation is associated with important debilitating diseases and cancers.^1–7^

Despite significant progresses in understanding the stepwise assembly of the spliceosome, the molecular mechanisms by which spliceosomal proteins interact together and mediate the ordered rearrangements within the spliceosome remain elusive. The multiple and dynamic protein-protein and protein-RNA interactions implicated in the assembly of the spliceosome machinery can be made possible by the large abundance of intrinsically disordered proteins (IDPs).

IDPs are characterised by the lack of stable secondary and three-dimensional structures. Due to their high conformational flexibility, IDPs are able to interact specifically, but transiently or weakly, with multiple protein partners, hence acting as hubs in protein-protein interaction (PPI) networks and molecular scaffolds that drive the formation of complex cellular machinery and membrane-less organelles (MLOs), such as Cajal bodies (CBs), nucleoli, or nuclear speckles, driven by liquid-liquid phase separation (LLPS).^8^

Recently, we demonstrated unrecognized functions for the G-Patch protein TFIP11 in the regulation of human spliceosome assembly and splicing efficiency.^9^ We observed that TFIP11 is located in multiple membrane-less organelles (MLOs) in which the snRNPs biogenesis and maturation take place, such as nuclear speckles, Cajal bodies (CBs) and nucleoli. Interestingly, we found that TFIP11 interacts with multiple key U4/U6.U5 tri-snRNP-specific proteins and we demonstrated that TFIP11 knockdown alters the assembly and stability of the U4/U6.U5 tri-snRNP complex, blocking the spliceosome in a B-like conformation. Consistent with our data, several other studies reported that TFIP11 is recruited during B complex formation and stably integrated at this stage, suggesting that TFIP11 is a crucial actor during the assembly and activation of the spliceosome machinery.^10^

It is well known that structural information can provide key insights into protein function. A deeper characterisation of TFIP11 structure might therefore provide a better understanding of its function and mode of action. No structural data for TFIP11 are available so far. However, *in silico* bioinformatic prediction suggested the presence of three intrinsically disordered regions (IDRs) in the whole TFIP11 sequence.^9^ The IDR#1 (residues 1-150) and IDR#2 (residues 175-250) surround the G-Patch domain (residues 148–193). The IDR#3 (residues 720-770) overlaps the nuclear speckles targeting site (NSTS) at the C-terminal extremity. The G-Patch domain constitutes a well-conserved glycine-rich domain present in other eukaryotic RNA-processing proteins such as PINX1^11^, SON^12^, GPKOW^13^, etc., and involved in protein-protein and protein-nucleic acid interactions.^14,15^ It is well-known that the G-Patch domain of TFIP11 interacts with and activates the ATP-dependent RNA helicase DHX15.^16^ Furthermore, the two predicted N-terminal IDRs in TFIP11 are also critical for interaction with coilin, the scaffold protein of CBs.^9^

In addition to IDRs, five low complexity domains (LCDs), LCD1 (residues 5–21), LCD2 (residues 82–96), LCD3 (residues 210–216), LCD4 (residues 226–240), and LCD5 (residues 291–306) are also predicted within the N-terminal region of TFIP11.^9^ LCDs are composed of a limited variety of amino acids and have distinct physico-chemical properties depending on the type of amino acid(s) each LCD is enriched with.^17^ LCDs are mainly found in RNA- and DNA-binding proteins, such as the EWS^18^ and TAR^19^ proteins, where they enforce gene regulation and functioning through the formation of dynamic complexes in MLOs such as CBs.^20^ Indeed, the LCDs can drive the formation of extended biomolecular condensates and MLOs by promoting LLPS.^21,22^ Together, these evidences strongly suggest that the N-terminal region of TFIP11, including the two IDRs and the G-Patch domain, play a central role in promoting not only the interactions with spliceosomal proteins but also with CBs scaffold protein, such as coilin, and are therefore of crucial importance to TFIP11 function.

By combining spectroscopic and scattering technics, as well as all-atom molecular dynamics (MD) simulations and disorder prediction algorithms, we provide here an additional and in-depth insight on the N-terminal (N-TER) TFIP11 structural and conformational properties. Given that LLPS formation and IDP conformations are salt-dependent^23,24^, the influence of salt concentration in shaping the TFIP11 N-TER conformational ensemble has been particularly investigated.

## 2. Materials and methods

### 2.1 Sequence-based bioinformatics prediction

Disorder-associated properties have been predicted from the known amino acid sequence of the TFIP11 (Uniprot ID: Q9UBB9) N-TER involving residue 1 to 350. Overall, the per-residue percentage of intrinsic disorder (PPID) has been determined considering the average of 20 disorder predictors available online. This includes the Predictors of Natural Disordered Regions (PONDR) series of algorithms VL-XT, XL1-XT, CAN-XT, VL3, and VSL2^25–27;^ Prediction of Intrinsically Unstructured Proteins (IUPred3) for short and long disordered segments^28^; Protein DisOrder prediction System (PrDOS)^29^; ESpritz algorithms respectively trained on X-ray, Disprot, and NMR datasets;^30^ non-evolutionary and evolutionary-based Prediction of Order and Disorder by evaluation of NMR data (ODiNPred) ^31^; deep-learning-based predictor metapredict (v2.2)^32^; NORSnet, Ucon, and MetaDisorder MD algorithms on the PredictProtein webserver^33^; NetSurfP-3.0^34^; putative function and linker-based Disorder Prediction using deep neural network (flDPnn)^35^; and finally DisoMine from Bio2Byte tools^36^. Disorder profiles were generated with the Rapid Intrinsic Disorder Analysis Online (RIDAO) tool^37^, comprising PONDR and IUPred predictions. On the disorder plot, residues with a score above the 0.5 threshold are considered as disordered, while flexible segments typically exhibit scores between 0.2 and 0.5.

The disorder propensity and the conformational ensemble of TFIP11 N-TER was further examined by cumulative distribution function (CDF) and charge-hydropathy (CH) plots obtained from the PONDR server^38^, as well as by the Das-Pappu phase diagram with the Classification of Intrinsically Disordered Ensemble Regions (CIDER)^39^. Namely, κ and Ω patterning parameters were also extracted from the CIDER analysis. The κ parameter describes the extent of charged amino acid mixing in a given sequence, while Ω describes the distribution of charged and proline residues with respect to all other residues. Presence and localisation of molecular recognition features (MoRFs) along the TFIP11 N-TER sequence have also been computationally identified with the MoRFchibi tool.^40^

### 2.2 Overexpression and purification of TFIP11 N-TER

TFIP11 N-TER recombinant protein was overexpressed with a 6xHis-tag at its N-terminus using a pET-like vector in *E. coli* BL21 (DE3) cells. Transformed bacterial strains were precultured with 0.36 mM ampicillin for 16 h at 37 °C, in 20 g/L lysogeny Lennox broth (LB). From 10.0 mL of preculture, strains were cultured with 0.14 mM ampicillin at 37 °C in 20 g/L LB, until the 600 nm-optical density reached 0.5–0.8. Cultures were then induced with 0.5 mM isopropyl β-D-1-thio-galactopyranoside (IPTG) at 18 °C for 18 h, and centrifuged. The resulted pellets were then stored at −20 °C. The pellets were then suspended in the lysis buffer (Tris-HCl 20 mM pH 7.4, 1% Triton X-100, 10% glycerol, 10µL/mL of Halt^TM^ Protease Inhibitor Cocktail (100X) (ThermoFisher), 1mg/mL lysozyme), and sonicated in an ice-water bath. Supernatants were discarded after centrifugation and inclusion bodies were solubilized in 25 mL buffer (Tris-HCl 20 mM pH 8, 6 M urea) at 8°C during 18 h.

The solution was filtered through 0.45 µm filter and purified on an Äkta Purifier fast protein liquid chromatography using Immobilized Metal Affinity Chromatography (IMAC). With the binding buffer (Tris-HCl 20 mM pH 8, 0.1 M NaCl, 6 M urea), His-TFIP11 N-TER fusion protein was bound on a 5 mL HisTrap^TM^ FF pre-packed column (Cytiva). The His-tagged protein was then eluted with elution buffer (Tris-HCl 20 mM pH 8, 0.1 M NaCl, 6 M urea, 0.5 M imidazole). Eluted fractions were gathered and further purified by size exclusion chromatography. A HiPrep^TM^ 16/60 Sephacryl S-200 HR (Cytiva) column was equilibrated (Tris-HCl 20 mM pH 8, 0.1 M NaCl, 6 M urea), then 2 mL of the IMAC purified protein were injected in the column and eluted using a flow rate of 0.5 mL/min.

The purity of the elution fractions was checked by sodium dodecyl sulphate-polyacrylamide gel electrophoresis (SDS-PAGE) and protein bands were revealed using Quick Coomassie Stain (CliniSciences). The protein sample is divided in two parts and transferred into a Tris-HCl 20 mM pH 7.4 buffer with 0 and 200 mM NaCl *via* a desalting column. *In vitro* measurements were performed immediately after the buffer exchange. The protein concentration was first determined by absorbance at 280 nm in the urea containing buffer since it prevents protein precipitation and avoids the misestimation of protein concentration caused by the effect of turbidity on the measured absorbance. The concentration of the protein is calculated from the one determined in urea with a dilution factor of 1.4 (due to buffer exchange). The concentration is estimated at 0.43 mg/mL (10 µM).

### 2.3 UV-visible spectroscopy

UV-visible spectra (200-600 nm) were recorded with a VWR® UV-6300PC UV-Visible spectrophotometer at 20 °C in Tris-HCl 20 mM pH 7.4, 0-200 mM NaCl using a 10 mm pathlength quartz QS cell (Hellma). Sample concentration was 0.43 mg/mL (10µM).

### 2.4 Fluorescence spectroscopy

Intrinsic tryptophan fluorescence (ITF) spectrum was recorded from the excitation wavelength of 295 nm up to 600 nm by 1.0 nm increment with a Shimadzu RF-6000 spectro fluorophotometer at 20 °C in Tris-HCl 20 mM pH 7.4, 0-200 mM NaCl, using a 10 mm pathlength quartz QS cell (Hellma), and an emission and excitation slit width (sw) of 5 nm.

### 2.5 Far-UV circular dichroism spectroscopy (far-UV CD)

Far-UV CD spectra (190–260 nm) were recorded with a MOS-500 spectropolarimeter at 20 °C in Tris-HCl 20 mM pH 7.4, 0-200 mM NaCl, using a 1 mm pathlength quartz Suprasil cell (Hellma). Four scans (15 nm/min, 2 nm bandwidth, 0.5 nm data pitch, and 2 s digital integration time) were averaged, baselines were subtracted, and corrected spectra were smoothed. Data are presented as the mean ellipticity ([Θ]ME), calculated as follows: [Θ]ME = (M.θ)/(10.C.l), where M is the molecular mass (Da), θ the ellipticity (mdeg), C the protein concentration (mg/mL), and l is the cell pathlength (cm). Sample concentration was 0.43 mg/mL (10 µM).

### 2.6 Dynamic light scattering (DLS)

DLS measurements were carried out at 20°C with Horiba Zetasizer SZ-100 nanoparticles analyser (detector at 90°). Protein samples were passed through a 0.2 µm PES filer before analysis. The auto-correlation function was successfully fitted 15 times per analyses. The results are expressed as the mean hydrodynamical diameter, 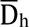 (nm).

### 2.7 Molecular dynamics (MD) simulations

All atoms equilibrium MD simulations of the TFIP11 N-TER, in different environmental conditions, have been performed in three independent replicates (triplicates) using the GROMACS 2020.5 suite.^41^ All the simulations have used the relaxed first-ranked AlphaFold2-generated model of TFIP11 as the initial structure (**Fig. S1**). The starting protein system was centred in a truncated octahedron water box including a 10 Å buffer from each edge and has been neutralised with the required minimal amount of Na^+^ cations. In a second system a NaCl concentration of 200 mM has been enforced, using the same protocol. The initial systems have been obtained using the *tleap* module of AmberTools. The AMBER ff14SB^42^ force field has been used to model the protein, while water has been described with the OPC model^43^. To better account for the TFIP11 disordered character, grid-based energy correction maps (CMAP) method, available for the amber ff14SB force fields, have been applied to the protein IDRs and the flexible segments, *i.e.* residues 1-24, 36-44, 57-143, and 166-337^44^. Hydrogen mass repartitioning (HMR)^45^, redistributing the mass of non-solvent hydrogen atoms has been consistently used to allow, in combination with Rattle and Shake, the use of a time step of 4 fs for the integration on the Newton equations of motion. HMR was enforced by modifying the initial topology with the *ParmEd* package available in Amber Tools.

Prior to the MD, the system has been minimised for 50000 steps, and subsequently thermalised and equilibrated. MD has been propagated in the isothermal and isobaric (NPT) ensemble. Constant temperature (300 K) and pressure (1 bar) have been maintained using a modified Berendsen thermostat and the Parrinello-Rahman barostat, respectively. Long-range electrostatic interactions have been calculated using the Particle Mesh Ewald (PME) summation^46^. Each MD replicate was propagated for 1 µs in periodic boundary conditions (PBC) in all three dimensions, energies and trajectories being recorded every 20 ps. Root-mean square deviation (RMSD) distribution, root-mean square fluctuation (RMSF), and radius of gyration (R_g_) time-series were calculated from the resulting trajectories, which have been visualised using PyMOL^47^ and VMD^48^.

## 3. Results and discussion

### 3.1 Sequence-based disorder-associated properties of TFIP11 N-TER

Given the complex behaviour of the N-TER domain of TFIP11, we aimed at refining in more details the bioinformatic predictions assessing and comparing complementary descriptors. Based on the average of 20 disorder predictors, the overall per-residue percentage of intrinsic disorder (PPID) globally amounts to 70% for the TFIP11 N-TER. The predicted PPID tendency is also corroborated by the content in order-promoting (OPRs) and disorder-promoting residues (DPRs). The latter considers the effects of short non-polar (glycine and alanine), charged (lysine, arginine, and glutamate), polar (serine and glutamine), and secondary structures breaking (proline) residues. As previously reported^9^, full-length TFIP11 has a higher proportion of DPRs (50.2%) compared to OPRs (34.2%). Yet, this tendency is further amplified for its N-TER domain, in which the amount of DPRs increases up to 59.4%, while the amount of OPRs is concomitantly decreased to 25.7%.

By using three independent algorithms (MFDp2 v2.0, PrDOS, and DISOPRED v3.1), two IDRs have been previously identified in TFIP11 N-TER at positions 1-150 and 175-250.^9^ We have refined the occurrence and location of IDRs with Rapid Intrinsic Disorder Analysis Online (RIDAO), generating and analysing the PONDR-FIT and IUPred predicted disorder profiles (**Fig. 1a**). Although some variability can be observed between the different predictors, even for the more disordered N-TER region, three main and discontinued IDRs clearly stand out showing a disorder score well above 0.5. The three IDRs are located at the N-TER region and are separated by shorter flexible segments for which the disorder score varies between 0.2 and 0.5. The first disordered region (IDR1) is comprised between residues 1 and 50 and is followed by two longer disordered regions comprising residues 85-145 (IDR2) and 190-265 (IDR3). Due to a much larger variability amongst the predictors for the residues comprised between the position 270 and 350, we restrain to assign this segment as an IDR. Nevertheless, scores close to the 0.5 threshold indicate that a large flexibility is still present in this region. Interestingly, the three IDRs have a high content of charged and hydrophilic residues (**Fig. S2**), a very common characteristic of IDP.^49^

**Fig. 1:**
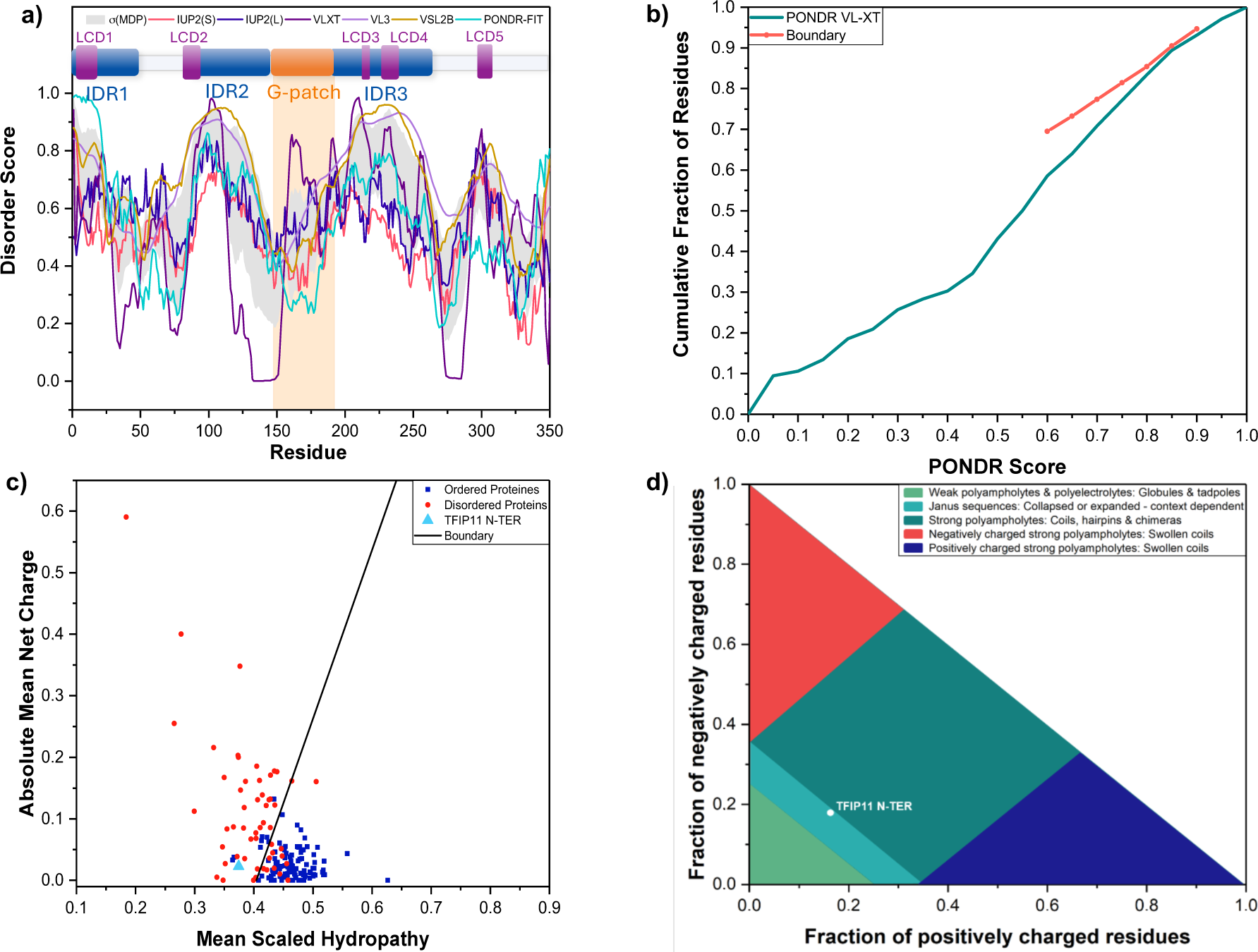
Predicted disorder-associated properties of TFIP11 N-TER. (A) PONDR and IUPred disorder profile, sequence and domains organisation of TFIP11 N-TER; MDP corresponds to the mean disorder profile. Every residue displaying a score above the 0.5 threshold is considered disordered, (B) CDF, (C) CH plots, and (D) Das-Pappu phase diagram.

TFIP11 N-TER disorder propensity was further investigated with the cumulative distribution function (CDF), as well as the charge-hydropathy (CH) plots (**Fig. 1b-c**).^50^ In the case of CDF plot, IDPs should consistently remain below the defined boundary, while ordered proteins (ORDPs) should overcome the threshold.^51^ The N-TER CDF profile is located slightly below the boundary, thus supporting its global disordered nature. Indeed, the presence of partially folded sub-structures linking the N-TER IDRs may shift the plot close to the CDF boundary. The difference in disorder content is far more noticeable on the CH plot, discriminating IDPs from ORDPs on the basis of their absolute mean net charge <R> with respect to their mean scaled hydropathy <H>.^52^ The TFIP11 N-TER region displays both low mean net charge and hydropathy and is found above the boundary defined by the equation <R> = 2.785<H> – 1.15, i.e. clearly laying in the IDP area, strongly supporting the assignation of the N-TER domain as a mainly disordered one.

The disorder-related conformational state of TFIP11 N-TER was further examined with the Das-Pappu phase diagram, as well as the κ and Ω parameters defined by the Classification of Intrinsically Disordered Ensemble Regions (CIDER) (**Fig. 1d**). Notably, the Das-Pappu phase diagram describes the conformational ensemble of a selected IDP from its fraction of positively against negatively charged residues.^53^ The TFIP11 N-TER region is found within the R2 region of the diagram, i.e. the region which comprises most of IDPs (∼40%). R2-associated IDPs are referred to as Janus sequences, reflecting their structural duality, namely the coexistence of ordered and disordered phases, also depending on their coupling with the environment. Indeed, R2-associated IDPs exist either as a collapsed ensemble (molten globule), an arrangement which is mostly driven by hydrophobic effect, or as extended conformations (random coil) driven by favourable protein chain solvation. Interestingly, the TFIP11 N-TER domain nearly reaches the R3 category (∼30% of IDPs), comprising strong polyampholytes with coiled or hairpin conformations.^54^

The analysis of the patterning parameters informs on the distribution of positively and negatively charged amino acids along the sequence (κ), as well as on the alternation of charged and proline residues (Ω). On a scale from 0 to 1, the closest the indexes are to 1, the more segregated the charged and/or the proline residues in the protein sequence are, thus resulting in more collapsed and compact conformers.^53,55^ TFIP11 N-TER has κ and Ω values of 0.24 and 0.14, respectively. κ and Ω values indicate the absence of extended patches of charged and proline residues, while their values remain sufficiently low to suggest that the N-TER region preferentially adopt more expanded conformations.

Molecular recognition features (MoRFs) are short disordered regions (10-70 residues) which can be found within a given protein sequence, and which are able to fold by binding to the interaction partners and, thus, enforce PPI in complexes involving IDPs.^56^ The number and the location of such segments in TFIP11 N-TER have been determined using the MoRFchibi tool. We have identified five MoRFs, which are consistently localised in the N-TER domain, and three of them are found in the defined IDRs (**Fig. 2**). Indeed, while IDR1 contains two MoRFs - MoRF1 (residues 1-15) and MoRF2 (residues 25-31) -, MoRF5 (residues 235-249) belongs to IDR3. Interestingly, two additional MoRFs, MoRF3 (residues 75-81) and MoRF4 (residues 175-180), are situated in the short flexible regions linking IDR1 to IDR2 and IDR2 to IDR3, respectively.

**Fig. 2:**
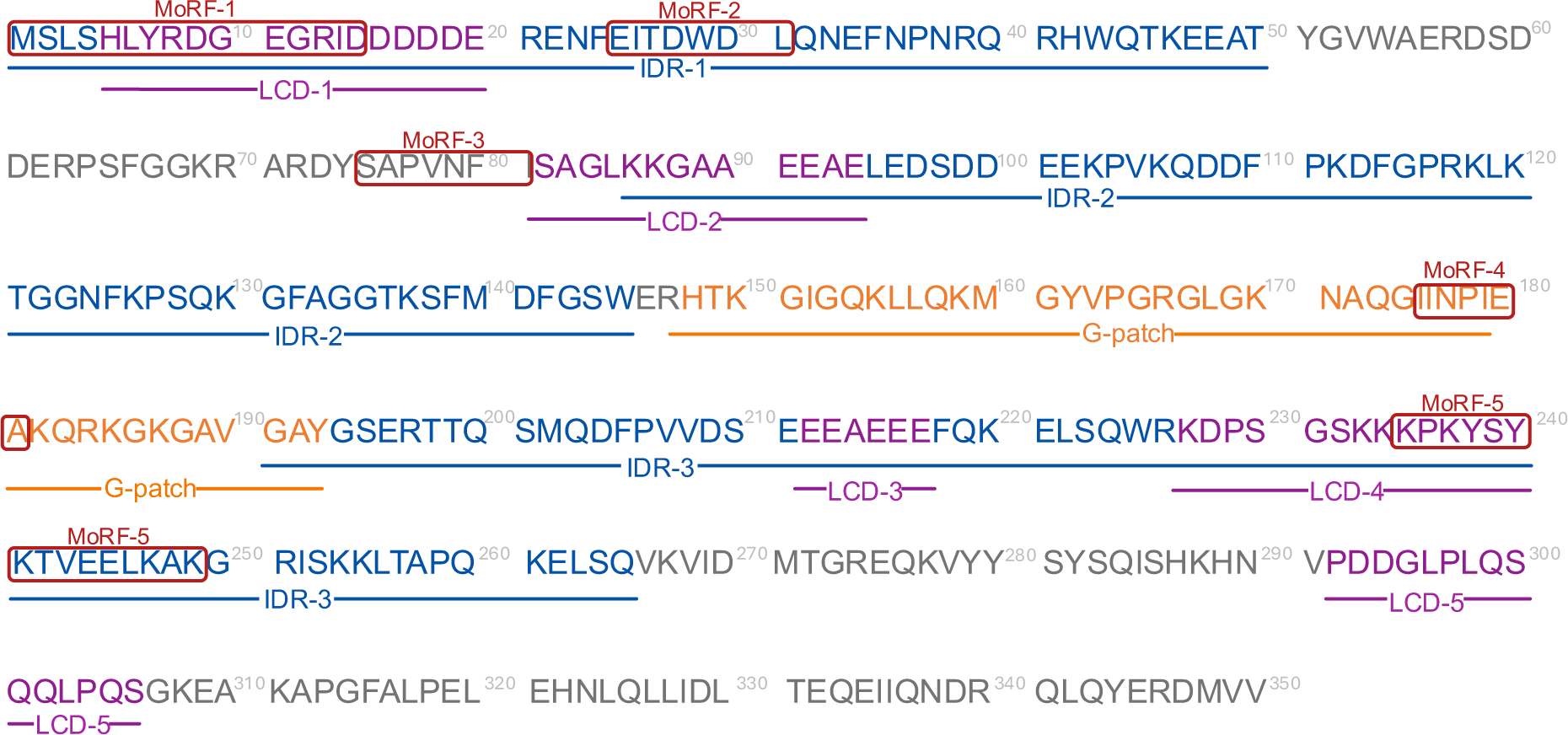
TFIP11 N-TER sequence with domain organisation (1-350 amino acid residues) comprising IDRs, LCDs, G-Patch, and MoRFs.

All together, these analyses suggest the classification of TFIP11 as an IDP and reveal that its main disorder area lies in its N-TER domain, as particularly evidenced by the disorder-associated predictions. Additionally, the presence of several MoRFs further supports the critical role of the N-TER region in TFIP11 functionality, probably in favouring PPI in molecular complexes.

### 3.2 In silico conformational properties of TFIP11 N-TER under two different ionic strength conditions

Since LLPS formation and IDP conformational changes are salt-dependent phenomena^23^, we have investigated by all-atom MD simulations the influence of salt concentration in shaping the conformational ensemble of the N-TER region of TFIP11.

First, the stability of the TFIP11 N-TER conformational ensembles at 0 and 200 mM NaCl concentration was compared, by calculating the backbone RMSD over the entire TFIP11 N-TER sequence for both conditions (**Fig. 3a; Fig. S3**). During the simulation, the overall fold of the protein in 0 mM NaCl remains stable despite its intrinsic disorder. However, after the first 400 ns of simulation, a slight increase in RMSD value is observed for the protein in 200 mM NaCl, suggesting that the presence of salt allows TFIP11 N-TER to adopt a slightly different conformational space. To assess the conformational variability along the MD simulations, the RMSD distribution of TFIP11 N-TER in 0 and 200 mM NaCl conditions are compared (**Fig. 3b**). Both systems display a broad distribution, which is coherent with the inherent flexibility of the protein. At 200 mM NaCl, the distribution is globally shifted towards higher values indicating that the structure deviates further from its starting conformation. These structural changes are highlighted by superimposing representative snapshots corresponding to the first 10 clusters in the two conditions (**Fig. 3b**). The protein at 200 mM NaCl has a fuzzier conformational landscape, a more globule-like shape, as well as an overall unstructured conformation as compared to the system at 0 mM NaCl. Furthermore, the 200 mM NaCl system also appears to present more accessible regions that might be involved in homotypic interactions, and hence should be more favourable to develop PPI networks.

**Fig. 3:**
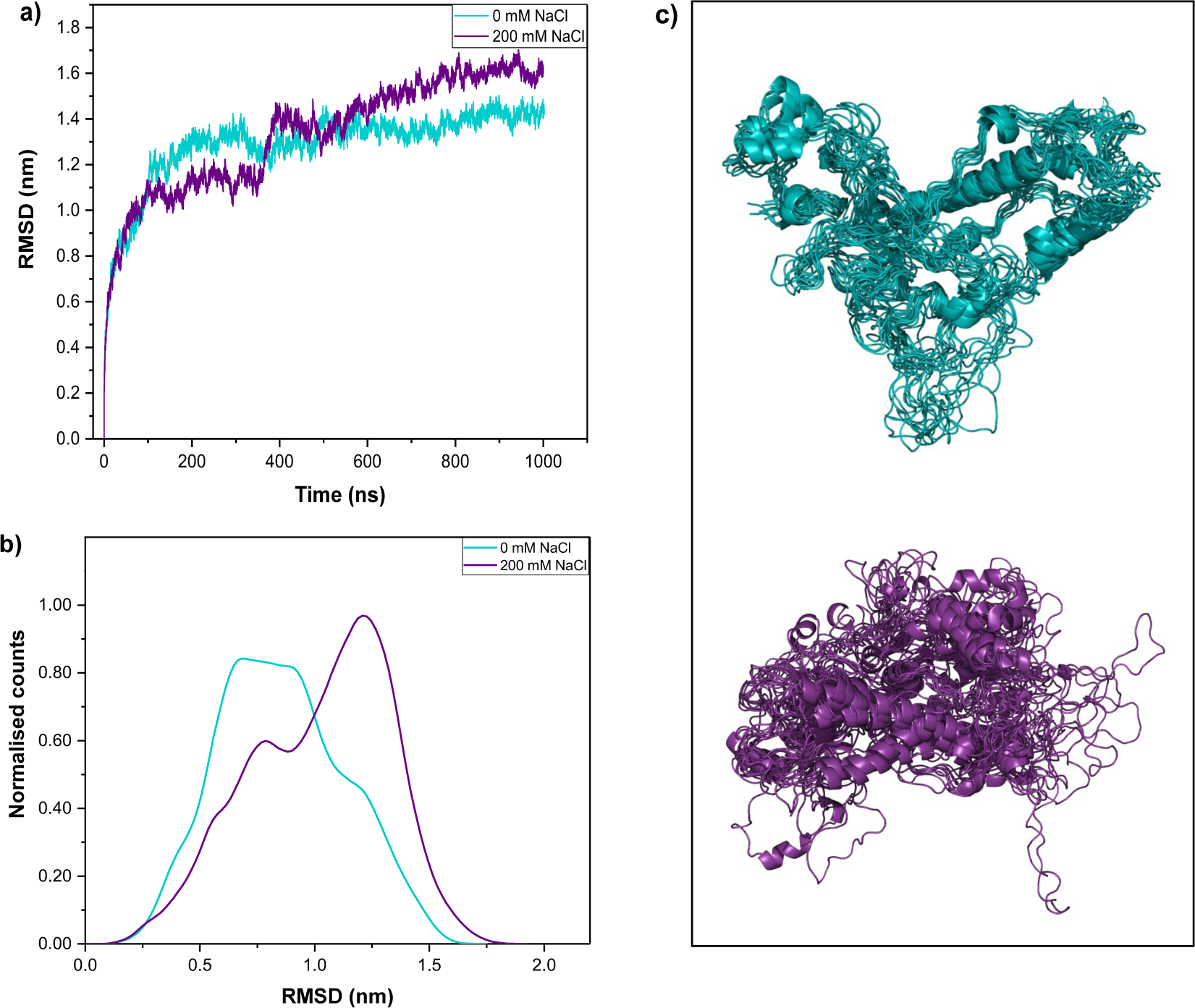
RMSD profile for TFIP11 N-TER. a) The RMSD of the backbone atoms from the equilibrated conformation (0 ns) is presented as a function of time, for 0 mM (blue) and 200 mM (violet) NaCl. b) RMSD distribution of TFIP11 N-TER. c) The first 10 clusters are superimposed for the system at 0 mM (blue) and 200 mM (violet) NaCl.

In addition, consistent with their flexible properties, the higher RMSF values for both conditions are unsurprisingly located in the IDRs and LCDs (**Fig. 4a; Fig. S4**). Interestingly, an important difference for the two salt concentrations is observed within LCD2, IDR2, G-Patch, and LCD5, which present almost systematically higher RMSF values in the 200 mM NaCl condition. These domains are comprised within the three regions (A, B, C) with the highest variation of RMSF (αRMSF) (**Fig. 4b**). The variation of RMSF is particularly high within the B and C regions containing IDR2, G-Patch, and LCD5, indicating a large backbone flexibility in these areas. Interestingly, G-Patch proteins are known to interact with helicases via their eponymous glycine-rich motifs. In the peculiar case of TFIP11, its G-Patch domain interacts with the RNA helicase DHX15.^57^ From a structural point of view, and given their sequence similarity, it is suggested that the G-Patch domain of TFIP11 could act in a similar way as the one of the ribosome biogenesis factor NKRF.^58^ Indeed, as determined in the crystal structure of the human helicase DHX15/Prp43 in complex with the G-Patch domain of NKRF, the latter acts like a flexible arm linking dynamic portions of DHX15 and tethering the two mobile parts of the protein together. The NKRF G-Patch motif is mostly unstructured and flexible, apart from a short N-terminal α-helix, which is in line with the structure of the B region in TFIP11 (**Fig. 4c**), and thus again confirms the role of structural flexibility in regulating TFP11 biological function. The structural changes between the two salt conditions highlight the environmental sensitivity of the conformation of the TFIP11 G-Patch, IDRs, and LCDs (**Fig. 4c**).

**Fig. 4.**
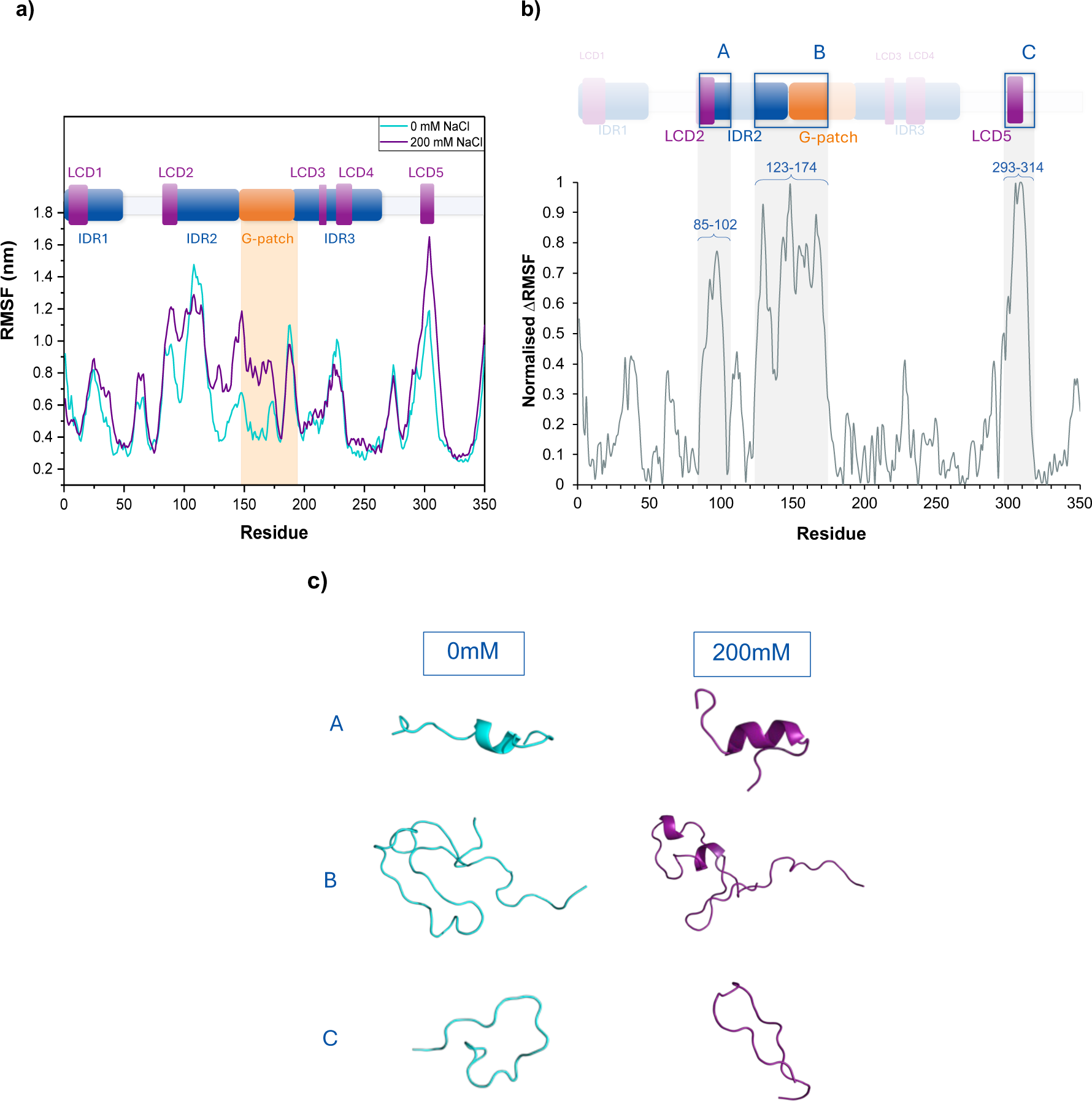
RMSF profile for TFIP11 N-TER. a) RMSF values of atomic positions computed for the backbone atoms as a function of residue number for 0 mM (blue) and 200 mM (violet) NaCl. b) Per residue normalised variation of RMSF between 0 and 200 mM NaCl condition, with the most impacted regions highlighted in gray and shown within the sequence domains. c) 3D structure of the TFIP11 N-TER regions (A, B, and C) impacted by salt concentration at 0 mM (blue) and 200 mM (violet) NaCl.

A further indication of the stability of the protein structure may be obtained by analysing the evolution of the radius of gyration (R_g_) (**Fig. 5a; Fig. S5**). We evidence that the increase in ionic strength does not correlate directly with significant changes in the R_g_ values. However, at 200 mM NaCl, the R_g_ presents a slight increase and variation going from an average value of 2.71 nm to 2.76 nm, correlating well with the variations in RMSD and RMSF. The average R_g_ values of TFIP11 N-TER are slightly higher than the ones for ordered protein sharing the same length, usually comprised around 2.0 ± 0.3 nm, further supporting its disordered features and its more extended conformation.^59^ Superimposed protein structures, extracted from the simulations for the two salt conditions (**Fig. 5b**), permit to visually appreciate the shape and conformational changes of the protein induced by the increase in the ionic strength, which, despite the modest effects on R_g,_ appear as substantial.

**Fig. 5:**
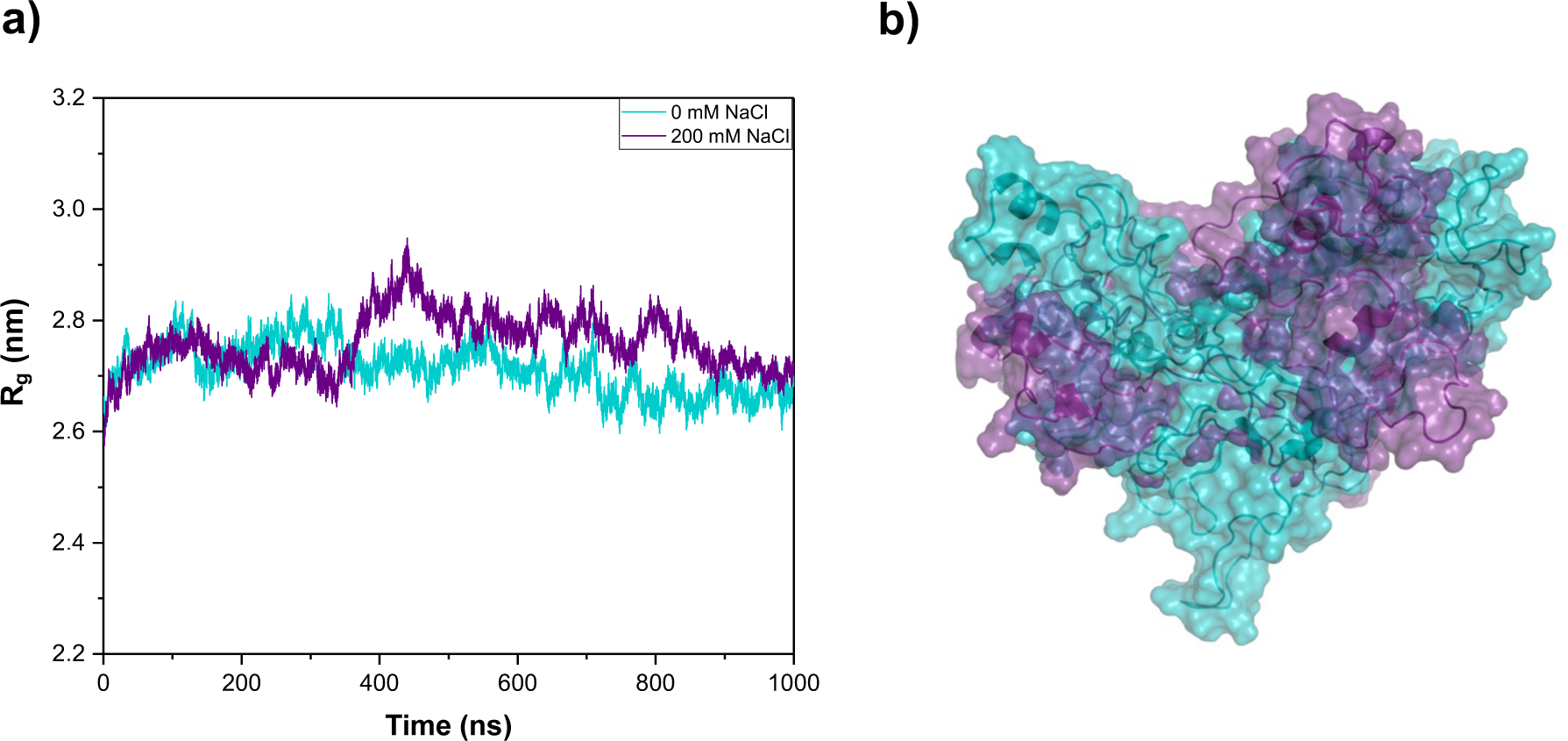
a) Radius of gyration (R_g_) versus time plot during the 1000 ns MD simulation for TFIP11 N-TER. The R_g_ values for 0 mM (blue) and 200 mM (violet) NaCl are shown. b) Surface representation of the superimposed first clusters for the system at 0 mM (blue) and 200 mM (violet) NaCl.

The intensity of the effects of salt concentration in shaping IDP conformation depends on the charge repartition within the protein, as well as on the salt concentration gradient.^24,60^ Generally, at low salt concentration, the changes in conformation are primarily driven by the possibility to maximise the screening of unfavourable electrostatic interactions. Indeed, polyampholyte IDPs, which is the case of TFIP11 N-TER (**Fig. 1d**), have been shown to undergo conformation expansion, that may lead to an increase of 5 Å in the R_g_ when going from 0 to 1 M salt concentration.^61^ Interestingly, such expanded conformation allows homotypic IDP-IDP interactions that further drive LLPS formation. This is the case of the Tau protein, a polyampholyte IDP involved in Alzheimer’s disease, the conformation of which extends within LLPS droplets.^62^ The weak, but significant, salt concentration-dependent increase in the R_g_ of TFIP11 N-TER is consistent with these observations. This leads to the hypothesis that TFIP11 could form LLPS under certain conditions, a characteristic which may be related to its biological functions inside MLOs.

To obtain a deeper understanding of the structural changes induced by salt concentration, the timeline analysis of the secondary structure content of the TFIP11 N-TER was performed (**Fig. 6**). Some structural features in the C-terminal end of the N-TER region (residues 209-225, 243-249, and 318-345), such as the α-helices, appears highly stable and persistent all along the MD simulation, and rather independent on salt conditions. This observation is also coherent with the low flexibility observed for these regions in the RMSF analysis, and by the fact that these motifs have been correctly predicted by AlphaFold (**Fig. S1**). Interestingly, this secondary structure stability also involves structural motifs, once more in the form of α-helices, that are embedded in the IDR1 and IDR3 regions. Yet, many regions (residues 3-7, 28-35, 46-51, 89-95, 154-160, 261-265, and 282-286) contain transient helices that vary between α and 3_10_ conformations, highlighting the dynamic nature of the TFIP11 N-TER. However, another stable structural motif involves the extended ß-sheet downstream of IDR3 (residues 268-270 and 277-279), which varies only slightly with salt conditions.

**Fig. 6:**
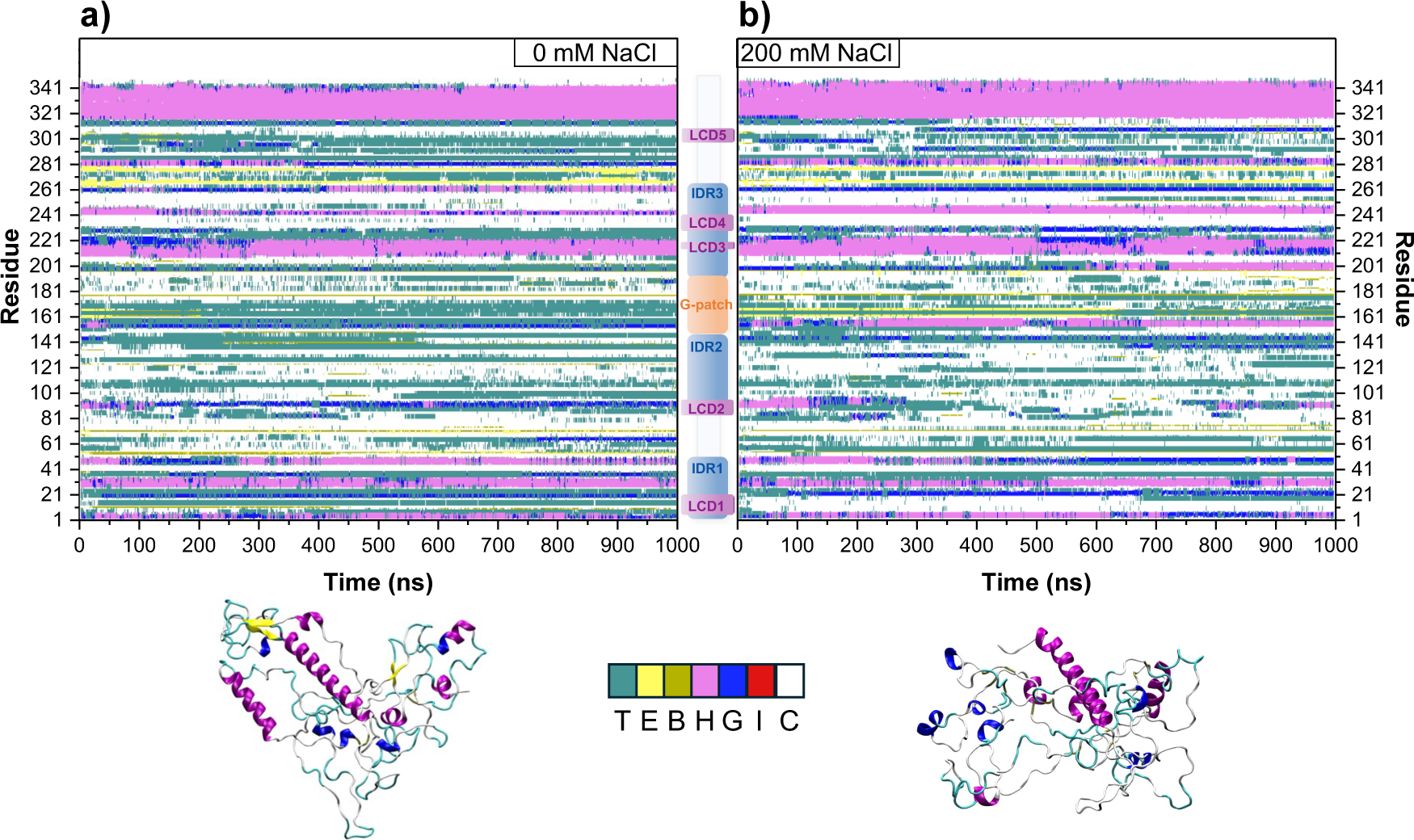
Secondary structure timeline analysis as computed by the timeline plugin contained in VMD. In the graphic, the extended β-sheet components (E) and isolated bridges(B) are represented in yellow and dark yellow, respectively; turns (T) are in teal; degrees of helix are α-helix (H) in pink, 3-10 helix (G) in blue, and π-helix (I) in red; random coil (C) is in white. 3D structure of the first cluster for both salt conditions at (a) 0 mM and (b) 200 mM NaCl are represented with the aforementioned colour-coded secondary structures.

Yet, a significant increase in random coil content at higher ionic strength is observed in every LCD (except for the shortest LCD3), the regions containing IDR2, IDR3, and the G-Patch, which is also accompanied by a more global modification in the secondary structure composition. In the case of LCDs, turns are the most commonly disrupted structural motifs in favour of random coil, with the greatest change observed within LCD1. The latter differs from the other similar sub-domains due to its high composition of consecutive negatively charged residues (HLYRDGEGRIDDDDDE, residues 5-20), which could be preferentially involved in electrostatic interaction with the environmental salt. The conformations corresponding to the first cluster show that the modification of turns into random coil participates to the expansion of the system in 200mM NaCl, specifically underlying the increased accessibility of LCDs and the G-Patch. This is coherent with previous observations reported in the literature concerning the role played by LCDs in driving LLPS.^63^

Coherently with the observations from the RMSF analysis, the G-Patch domain also appears highly modified by increasing the salt concentration. In particular, we observe a strong decrease of turns in favour of extended conformations. Interestingly, the formation of an α-helical motif at 200 mM NaCl between the two first conserved glycine residues in the G-Patch, similarly to the behaviour observed for NKRF, highlights the sequence and conformational similarity between the two proteins.^64^ It is also important to underline that high ionic strength conditions appear to favour the presence of random coil arrangement in the IDR2 domain, which is also one of the protein regions most affected by the salt concentration. These data suggest that IDR2 and the G-Patch regions are interdependently involved in conformational changes.

### 3.3 In vitro characterisation of TFIP11 N-TER under two ionic strength conditions

To complement and support the analyses obtained from the disorder prediction algorithms and MD simulations, the purified recombinant TFIP11 N-TER (**Fig. S6**) was characterised by spectroscopic (CD and ITF) and light scattering (DLS) techniques in two ionic strength conditions (0 and 200 mM NaCl).

First, the far-UV CD signature (**Fig. 7**), providing secondary structure information, of the protein in 200 mM NaCl is characteristic of random coil with a strong negative band at 205 nm. The very weak positive band between 195 and 200 nm further demonstrates for the first time at experimental level that the human TFIP11 N-TER is mainly disordered. The broad shoulder between 220 and 225 nm nevertheless indicates the presence of some folded elements which can be likely attributed to ⍺-helix and/or anti-parallel ß-sheet. This is coherent with the *in silico* secondary structure analyses showing the presence of some ⍺-helices and ß-sheet structures (**Fig. 6**). In 0 mM NaCl, a significant change is observed in the CD spectrum revealing two broad negative bands at 220 and 226 nm, as well as a broad positive band centred at 199 nm, indicating a significant decrease in the disorder content to give a predominance of ß-sheets. This observation not only corroborates the MD simulations, showing an increase of disorder with the ionic strength, but also the disorder prediction algorithms, revealing a structural duality with the coexistence of ordered and disordered structures depending on the environment.

**Fig. 7:**
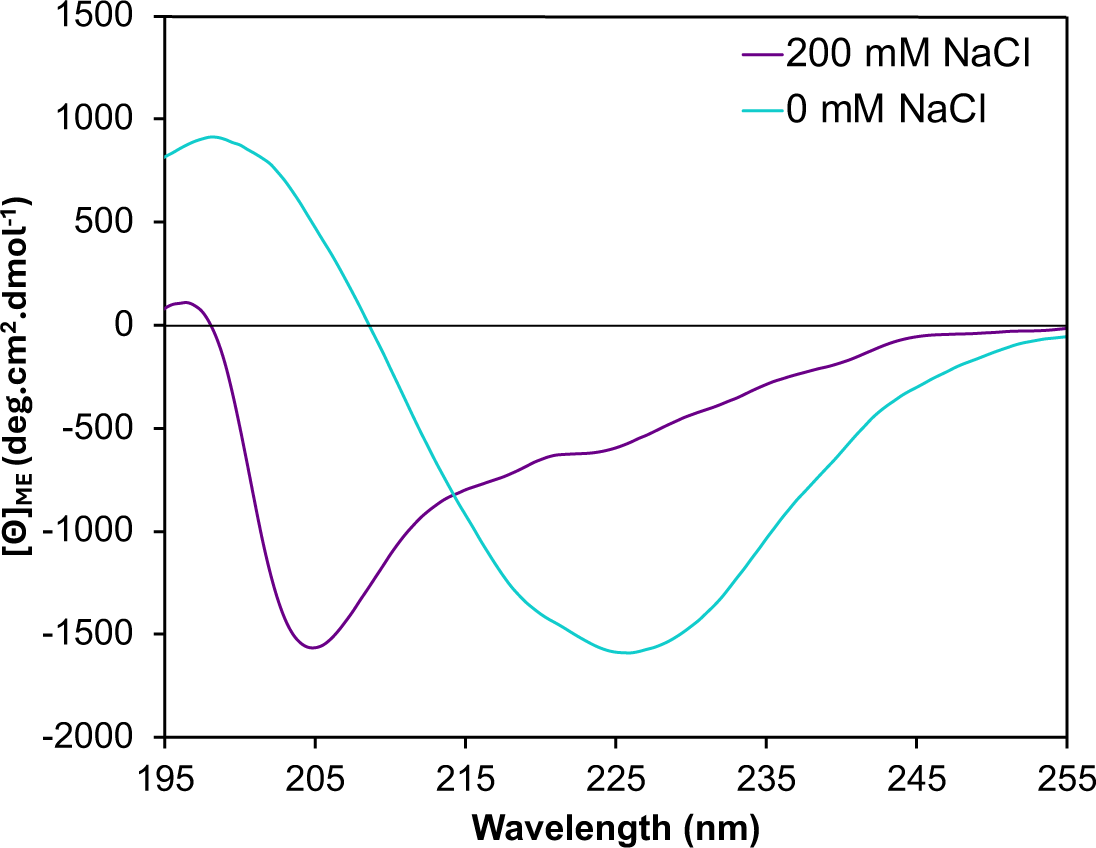
Far-UV CD spectra of His-TFIP11 N-TER in Tris-HCl 20 mM pH 7.4 with 0 mM NaCl (blue) and 200 mM NaCl (violet) at 20 °C. The protein concentration is 0.43 mg/mL (10 µM).

The presence of salt might favour electrostatic interactions between the numerous polar and charged residues contained in the IDP with ions in solution, leading to a more disordered and extended conformation, whereas the absence of salt might favour inter- and/or intramolecular interactions leading to more stable secondary structures. Interestingly, the CD footprint observed at 0 mM NaCl shows an enrichment in ß-sheets, suggesting an amyloid-like fibrillation of the protein, as already observed for various IDPs which are prone to aggregate.^65,66^ Further research on this protein will have to address this possibility.

Study of the intrinsic fluorescence properties of tryptophan (Trp) residues provides local conformational information about the protein.^67^ The ITF spectrum (λ_ex_ = 295 nm) (**Fig. 8a**) of the TFIP11 N-TER in 200 mM NaCl shows a maximum emission signal at 339 nm corresponding to Trp residues partially exposed to the solvent and/or polar amino acids. In 0 mM NaCl, a slight blue shift of the emission band (336 nm) is observed, revealing that Trp residues become less exposed due to a conformational change. ^67,68^ Interestingly, a second band appears at 470 nm, which is most likely associated to deep-blue autofluorescence (dbAF), an intrinsic fluorescence mainly found in proteins forming amyloid fibrils.^69^ This peculiar dbAF phenomenon is not exclusive to amyloid fibrils but can also be found in monomeric proteins. Nevertheless, the presence of such dbAF signal, in conjunction with the main ß-sheet CD signature, suggests that in the absence of salt, TFIP11 N-TER might tend to assemble into stable aggregates and adopt more ordered structures.

**Fig. 8:**
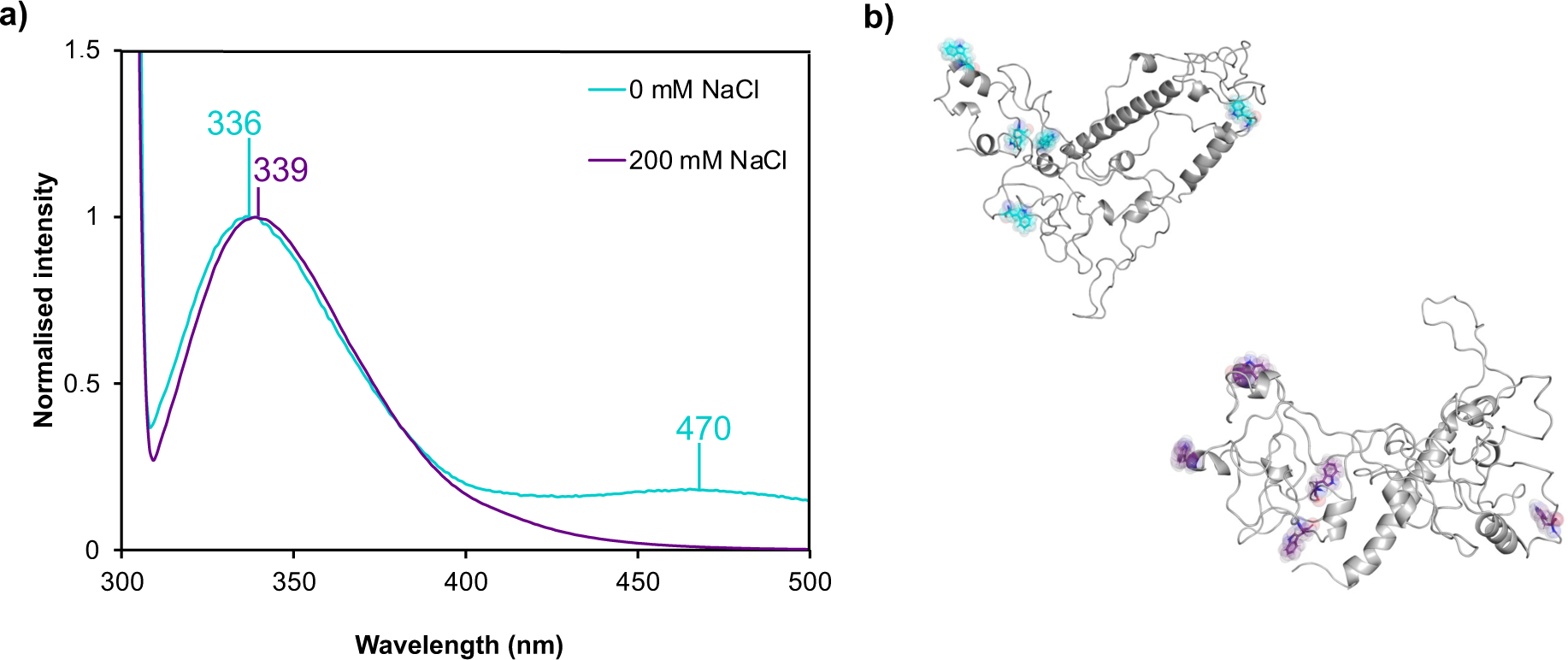
(a) Normalised ITF (λ_ex_ =295 nm, sw 5 nm) of His-TFIP11 N-TER in Tris-HCl 20 mM pH 7.4, 0 mM NaCl (blue) and 200 mM NaCl (violet) at 20 °C. (b) Snapshots of the first cluster of TFIP11 N-TER MD simulation with Trp residues shown in blue for the system in 0 mM NaCl and in violet for the system in 200 mM NaCl.

ITF data are in agreement with MD simulations, pointing out that the conformational ensemble of TFIP11 N-TER is more disordered in 200 mM than in 0 mM NaCl, due to the maximisation of electrostatic interactions with the ions in solution. This indicates a conformational dynamic where the state of Trp residues can vary from fully exposed to more buried depending on the environment. This is supported by the fact that four Trp residues are present in the IDRs (W29 and W43 in IDR1; W145 in IDR2; W225 in IDR3) making them more prone to changes of exposure in different environments. Furthermore, the highlighted Trp residues within TFIP11 N-TER conformation in 0 and 200 mM NaCl (**Fig. 8b**) permit to visually appreciate their position inside the protein and changes in their direct environment.

For a preliminary assessment of the assembled state of the protein as a function of the ionic strength, DLS measurements were performed. A 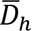 value of ∼199 nm and ∼136 nm was observed in 0 and 200 mM NaCl, respectively. These results first confirm the assembled state of TFIP11 N-TER in 0 mM NaCl. Unexpectedly, it seems also to reveal an assembled form in higher ionic strength. However, all our results would suggest two types of protein assemblies with either disordered or ordered containing structures. IDR containing proteins are known to form diverse types of assemblies that can either be in the form of reversible dynamic condensates or irreversible stable aggregates.^70^ Protein amino acid composition, concentration and physical-chemical properties of the environment promotes a certain type of assembly. Dynamic condensates are formed *via* LLPS and mainly found in MLOs such as nucleoli and Cajal bodies where TFIP11 was localised. Considering the disordered CD signature and the partially exposed ITF signal, the 200 mM salt-assemblies are hypothesised to be in a more extended and disordered conformation that might form biomolecular condensates driven by LLPS. The phase separation can then initiate protein aggregation, as it has been reported for Tau protein.^71^ On the other hand, in 0 mM NaCl, the protein assemblies might be of amyloid-like structure or another type of structured aggregates considering the predominant ß-sheet CD signature and the dbAF signal observed on ITF spectra.

Altogether, those experimental data clearly indicate that TFIP11 N-TER belongs to the class of polyampholyte IDPs. Indeed, TFIP11 N-TER shows a structural duality between disordered and ordered states, the prevalence of which relies on salt concentration, *i.e.* an increase in disorder with salinity. The different spectral signatures indicate that salt concentration induces different conformations that promote the formation of distinct types of protein assemblies. Although their correct identification requires further investigation, they have been reported in complex-forming and prone-to-aggregate IDPs^72,73^.

### 3.4 Role of putative post-translational modification sites in TFIP11 N-TER

IDPs are prone to post-translational modifications (PTMs) which can regulate their functions, namely, interaction with other protein partners, protein folding, etc. and, consequently, can change protein functions in various biological processes. PTMs of RNA-binding proteins (RBPs) can directly enhance or reduce their interactions with other proteins and/or RNA components, contributing to the formation of MLOs. TFIP11 is known to undergo extensive PTMs. In yeast, a lysine (K) residue at position 67 within the G-Patch domain of Ntr1/Spp382 (the yeast ortholog of human TFIP11 in yeast) is important for the binding of the protein to RNA.^74^ This K67 residue is equivalent to K residue at position 155 in the G-Patch of human TFIP11 and is highly conserved across species (**Fig. S7**), suggesting a crucial importance of this residue for the structure and function of TFIP11. In addition, this specific K155 is reported by the PhosphoSitePlus^75^ database as well as in the litterature^76^ as an acetylated residue, suggesting an acetylation-dependent regulation of TFIP11 function.

Interestingly, Tannukit S. *et al*. ^77^ reports that one conserved tyrosine residue (Y162) located in the G-Patch domain is phosphorylated and required for binding to nucleic acid.^78^ The presence of this phospho-Y162 in the G-Patch, predicted to contain two α-helices with four out of the six glycine residues located within an intervening loop, is consistent with the observation that phospho-Tyr is more often observed at ordered interfaces.^79,80^ In addition, a proteome-wide analysis of arginine monomethylation reveals that the arginine residue at position 166 (**R166**) located in TFIP11 G-Patch domain is mono-methylated and sensitive to PRMT1/4/5 inhibition.^81^ This R166 residue is adjacent to phospho-Y162 residue, which is consistent with Larsen S. C., *et al*. study^81^ reporting that arginine methylation sites regulated by PRMT1/4/5 are often found to be adjacent to phosphorylation sites. Moreover, this R166 is located between two neighbouring glycine residues (GRG), a preferential site for the PRMT5 enzyme^82^, a protein methyltransferase which has been detected in B spliceosomal complex.

All these observations lead us to speculate that PTMs in functional domains of TFIP11, such as the G-Patch, could be the molecular basis for binding to spliceosomal proteins and/or RNA substrates, and consequently contribute to the structural arrangement and activation of the spliceosome complex. To further support this assumption, the G-Patch domain was deeply examined. Interestingly, the three residues (Y162, K155, and R166) described above are located within the B region (**Fig. 4b**; **Fig. S8**), presenting a high variation of RMSF, indicating the large impact of salt concentration in this region containing both IDR2 and the G-Patch. For K155, the presence of salt modifies the interactions of such a positively charged residue, which previously interacted with two negatively charged residues (D34 and E47), to finally become totally accessible to the solvent, maximising electrostatic interactions with ions in solution (**Fig. 9**). This gain in accessibility may favour interactions with RNA and/or facilitate the addition of PTM, such as acetylation, to dynamically regulate RNA binding. The R166 residue also sees its environment being changed by the presence of salt. By switching from a π-cation interaction with residue W145 to an electrostatic interaction with residue D18, R166 participates in bringing the G-Patch and LCD1 (two types of domains found in RNA-processing proteins) closer together. Although residues K155 and R166 are found close to Y162, the latter is not significantly affected by the presence of salt, probably because its hydrophobic nature makes it less sensitive to salinity than charged residues. Nevertheless, addition of a negative charge to Y162 through phosphorylation could significantly change the conformation of the G-Patch and thus impact its function.

**Fig. 9:**
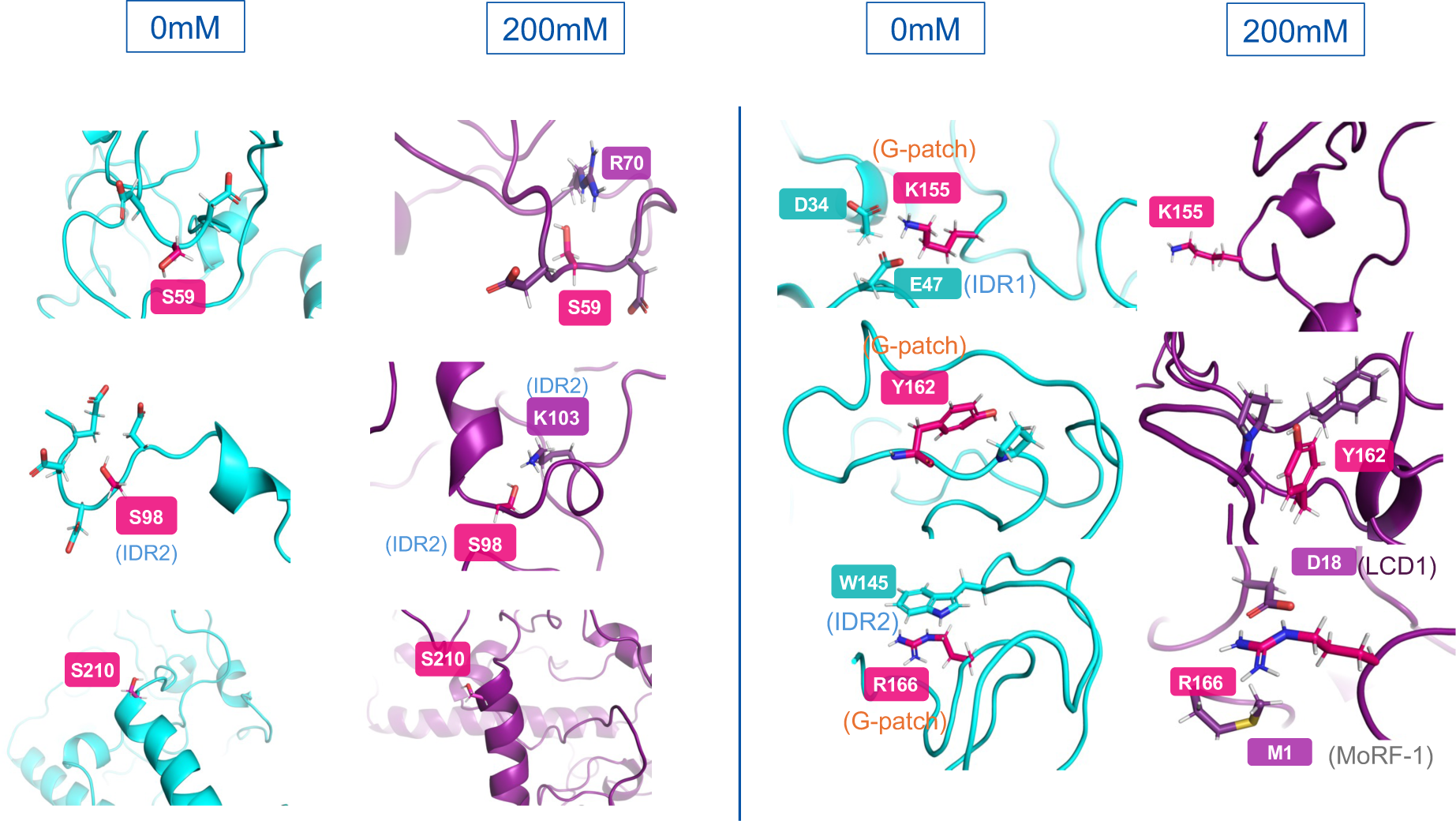
Putative PTM residues S59, S98, S210, K155, Y162, and R166 (in pink) with their modified environment and surrounding residues within TFIP11 N-TER MD predicted structure at 0 mM (in blue) and 200 mM (in violet) NaCl.

Another PTM frequently predicted in TFIP11 is the phosphorylation of multiple serine residues, such as S59, S98, and S210 (**Fig. S8**) (see PhosphositePlus database for all references for each residue). Therefore, we decided to compare the variation in structure and interactions of such residues according to salt concentration. S98 is located in the high ΔRMSF region A (**Fig. 4b**) in IDR2. Its interaction with K103, a charged residue within IDR2, is favoured by the presence of salt. A similar effect is observed for S59. At 200 mM NaCl, an electrostatic interaction with R70 is also promoted. The salt-induced proximity of these serine residues to positively charged residues shows that the charge of the system locally influences intramolecular interactions. This suggests that positively charged R or K residues, known to be involved in interactions with nucleic acids^83^, could be attracted by a local negative charge introduced by the addition of a phosphate group.^84^ This feature is frequently found in phosphorylated splicing factors, such as SF1^85^ and ASF/SF2, and is known as an “arginine claw” in reference to the cluster of arginine residues around the added phosphate group.

To further study PTMs, we investigated short linear motifs (SLiMs), linear protein interaction sites containing ligand motifs that mediate PPI and post-translational modification motifs, which are directly recognised and targeted for PTM by regulatory enzymes.^86^ The number and the location of predicted SliMs in TFIP11 N-TER have been determined using the Eukaryotic Linear Motif (ELM) resource prediction tool.^87^ The predicted SliMs are shown in **Fig. S9**. Only those contained in the flexible regions A, B, and C are listed. We focus on SliMs involved in the regulation of LLPS. For example, the SliM (residues 155-159) located within the G-Patch is predicted as a Ubiquitin carboxyl-terminal hydrolase 7 (USP7 CTD) domain binding motif. This type of hydrolases is responsible for the deubiquitination of target proteins. This is interesting given that K155 is also predicted by PhosphositePlus as being ubiquitinylated (as well as acetylated). A USB7 MATH domain binding motif is also predicted in LCD5 within region C (residues 297-301). Ubiquitination is known to modulate LLPS for many proteins such as Tau^88^, UBQLN2^89^, and HECT E3^90^. Depending on the protein, ubiquitination can promote or inhibit the formation of LLPS.^91^ Another interesting motif is the SLiM (residues 303-309) found in the C region comprising LCD5 and predicted as a phosphorylation site for casein kinase 2 (CK2). This kinase plays a key role in IDP folding.^92^ For example, the phosphorylation of FRMP by CK2 increases the density of negative charge in its IDRs and increases electrostatic interactions leading to LLPS.^93^ Studying the effect of phosphorylation by CK2 and other enzymes involved in the PTM of TFIP11 will provide a better understanding of the mechanisms regulating the LLPS phenomenon within the cell.

Further structural and functional studies on post-translationally modified and repositioned residues will allow to identify their role in the multiple biological functions of TFIP11.

## 4. Conclusions

The present study has provided a better understanding of the intrinsic and environment-dependent structural behaviour of the TFIP11 N-terminal domain. By combining disorder prediction algorithms, MD simulations, and spectroscopic techniques, we highlighted that the protein enters the definition of polyampholyte IDPs, involving a structural duality with the coexistence of ordered and disordered phases, depending on the environment. Our data emphasised that more flexible, extended, and unstructured conformations populated the high ionic strength condition. The regions which were mostly impacted by the increase in salt concentration were those mostly composed of charged and hydrophilic residues, *i.e.* LCD2, IDR2, G-Patch, and LCD5, coherently with their intrinsic flexible nature. The expanded conformation and more accessible domains (especially LCDs) within polyampholyte IDPs are known to allow homotypic IDP-IDP interactions that further drive LLPS formation and may consequently be strongly related to the biological role of TFIP11. Interestingly, we also showed that the putative PTM sites in the TFIP11 N-TER are present in the IDR2/G-Patch zone and are highly impacted by the environment.

Our experimental investigation has suggested the protein tendency to assemble in two very different forms: i) in low salt condition, as stable structured assemblies, and even potentially in amyloid(-like) fibrils, and ii) in high salt condition, as dynamic assemblies that may form biomolecular condensates, maintaining a high degree of disorder in the bound state. The condensate formation hypothesis, though requiring further investigations, is supported by the similarity in structural features of TFIP11 N-TER with polyampholyte IDPs undergoing LLPS such as the Tau protein.

In further prospects, our data consequently offer hints on the molecular basis leading to LLPS and hence MLO formation. This in turn will allow a better understanding of the spliceosome assembly process, as well as the key role of TFIP11 in promoting the complex and finely tuned regulations of different biological processes.

## CRediT authorship contribution statement

Blinera Juniku; Conceptualisation; Methodology; Validation; Investigation; Visualization; Writing – original draft; Writing - Review & Editing

Julien Mignon: Conceptualisation; Investigation; Writing - Review & Editing.

Rachel Carême: Validation; Investigation.

Alexia Genco: Review & Editing.

Anna Maria Obeid: Review & Editing.

Denis Mottet: Conceptualisation; Supervision; Writing - Review & Editing; Funding acquisition.

Antonio Monari: Supervision; Writing - Review & Editing.

Catherine Michaux: Conceptualisation; Methodology; Supervision; Writing - Review & Editing; Project administration; Funding acquisition.

## Funding and acknowledgements

B. J. and J. M. thank the Belgian National Fund for Scientific Research (F.R.S.-FNRS) for their FRIA (Fund for Research training in Industry and Agriculture) Doctoral grant and Research Fellow fellowship, respectively. A.M.O. and A.G are PhD students supported by the University of Liège (Fonds Spéciaux Recherche) and the Belgian National Fund for Scientific Research (F.R.S.-FNRS), respectively. The authors are appreciative to the PTCI high-performance computing resource and the Research Unit in Biology of Micro-organisms of the University of Namur. The present research benefited from computational resources provided by the Consortium des Équipements de Calcul Intensif (CÉCI), funded by the FNRS under grant n°2.5020.11 and by the Walloon Region, and made available on Lucia, the Tier-1 supercomputer of the Walloon Region, infrastructure funded by the Walloon Region under the grant agreement n°1910247. C. M. and D. M. also thank the FNRS for their Senior Research Associate position. A. M. thanks ANR and CGI for their financial support of this work through Labex SEAM ANR 11 LABEX 086, ANR 11 IDEX 05 02. The support of the IdEx “Université Paris 2019” ANR-18-IDEX-0001 is also acknowledged. B. J. thanks Younes Bourenane Cherif for his precious help in designing the figures.

## Appendix A. Supplemental data

Supplementary data to this article can be found online at https://github.com/blinerajuniku/Intrinsic-Disorder-and-Salt-dependent-Conformational-Changes-of-the-N-TER-TFIP11-Splicing-Factor-.git

## Notes

### Competing Interest Statement

The authors have declared no competing interest.

### Summary of Updates

The main points that have been added are: an experimental part (Point 3.3), and a discussion on the role of putative post-translational modifications sites in regard to molecular dynamics studies (point 3.4). In addition, one author was added (Rachel Careme).

