## supplementary information for "Intrinsic Disorder and Salt-dependent Conformational Changes of the N-terminal Region of TFIP11 Splicing Factor"

**AlphaFold predicted structure of TFIP11 N-TER**

The starting structure for MD simulation is the AlphaFold predicted structure which is represented with regions represented in different colours depending on the confidence score of the prediction. The regions highlighted in gray are the ones with high variation in RMSF (see Figure 4b in the main text).

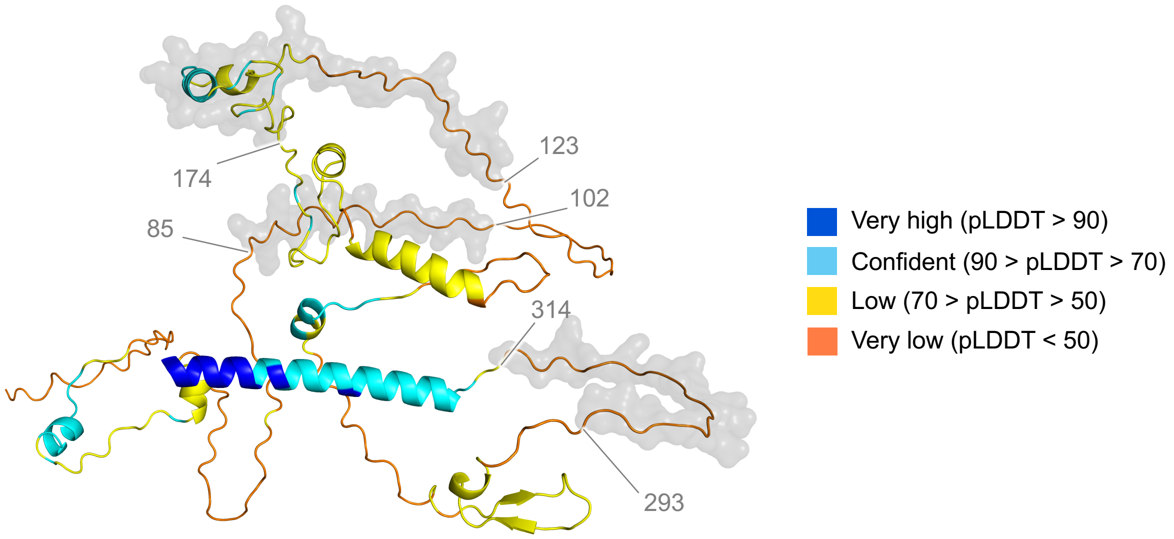


Figure S1: AlphaFold-predicted structure of TFIP11 N-TER.

**Intrinsically disordered regions (IDRs) composition in different amino acid types**

The composition in different types of amino acid within IDR-1, IDR-2, and IDR-3 is shown in different colors in the sequence and in percentage of each type in the circle charts.

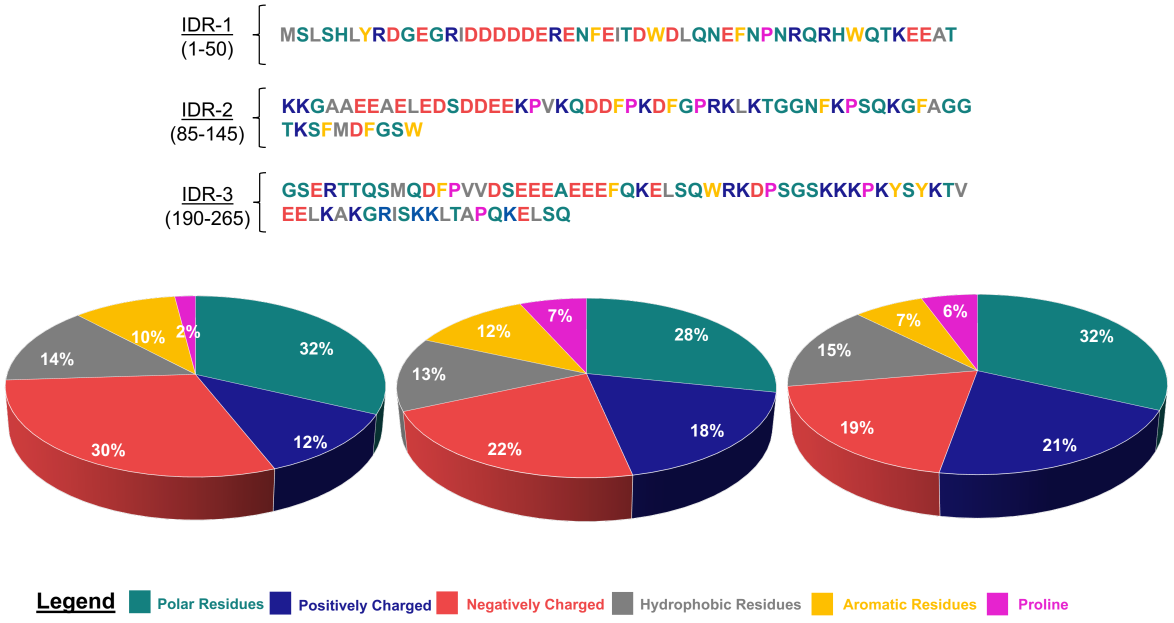


Figure S2: Percentage of polar, positively charged, negatively charged, hydrophobic, aromatic, and proline residues within IDR-1, IDR-2, and IDR-3 of TFIP11 N-TER.

**RMSD time series for all the MD simulations**

To evaluate when systems reach an equilibrium, we report the root mean square deviation (RMSD) for the systems containing TFIP11 N-TER in 0 and 200 mM NaCl. Three replicates are performed for both conditions.

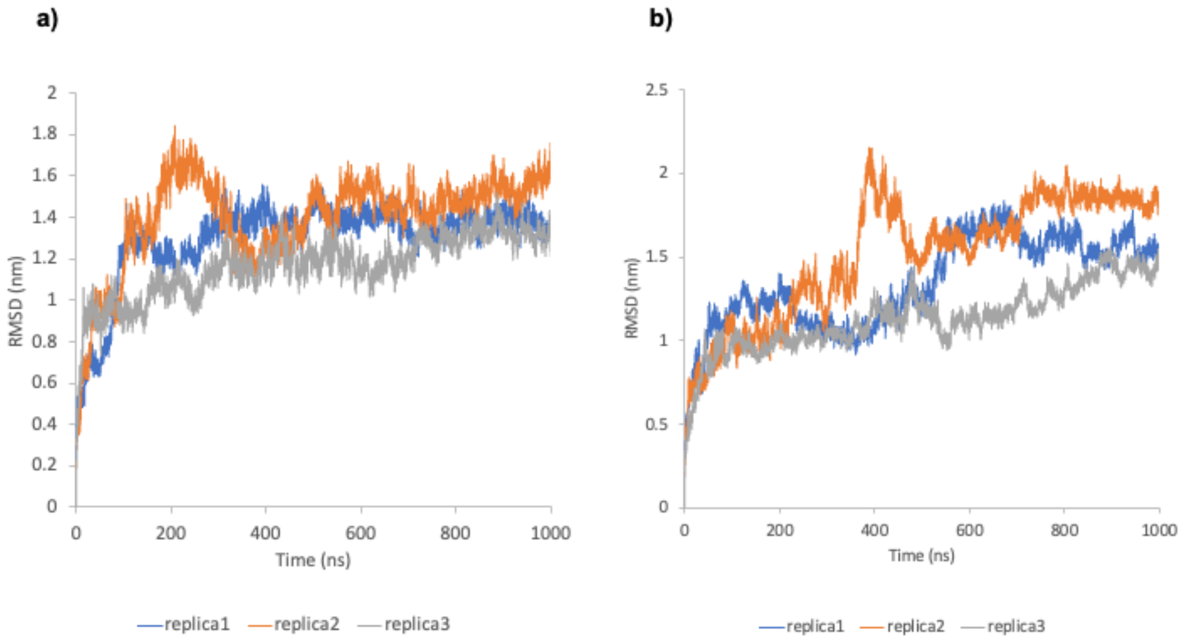


Figure S3: RMSD replicates for the system at a) 0 mM NaCl and b) 200 mM NaCl.

**RMSF time series for all the MD simulations**

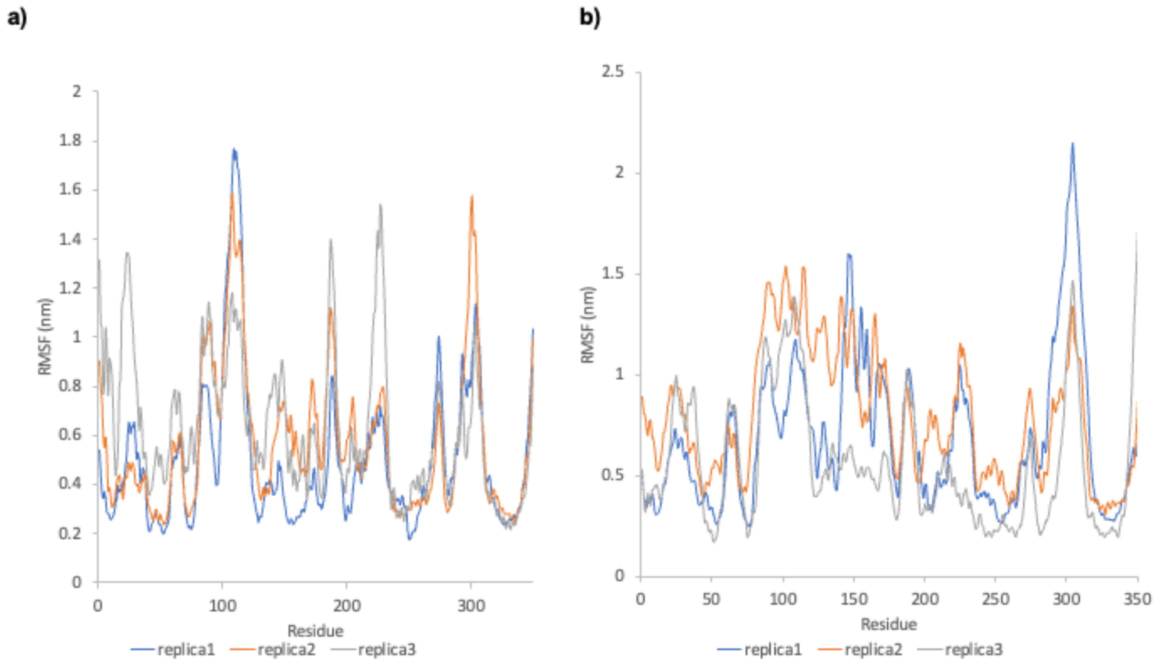


Figure S4: RMSF replicates for the system at a) 0 mM NaCl and b) 200 mM NaCl.

### $R_g$ time series for all the MD simulations

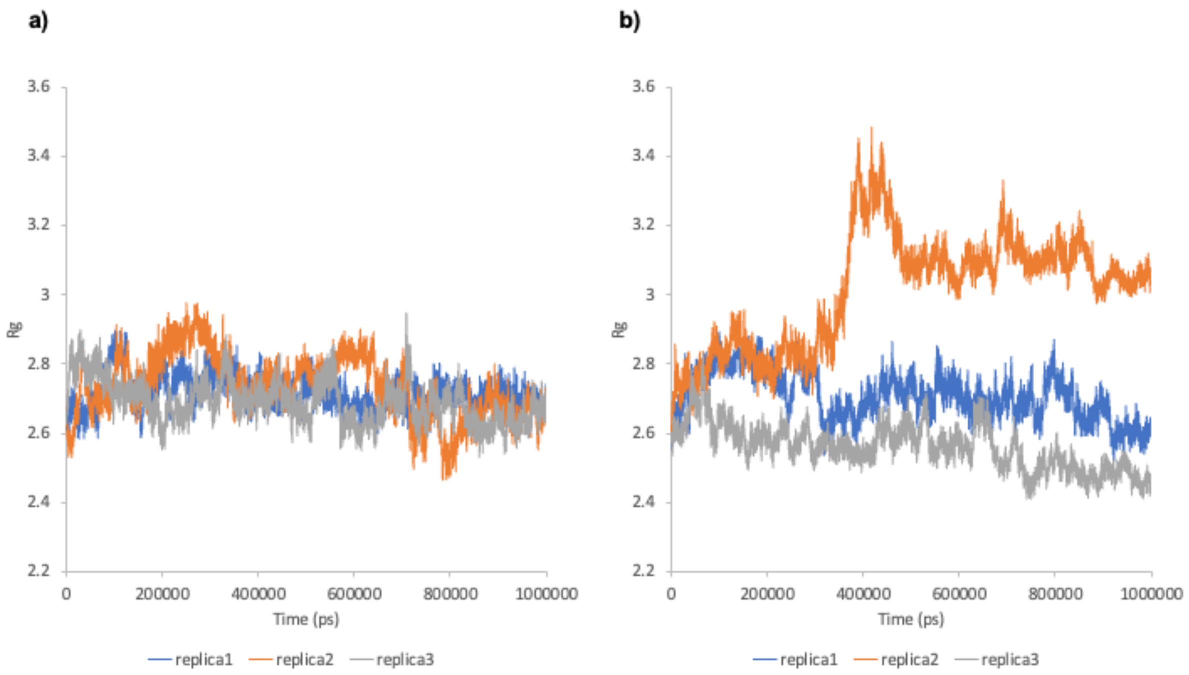


Figure S5: $R_g$ replicates for the system at a) 0 mM NaCl and b) 200 mM NaCl.

**TFIP11 N-TER purification**

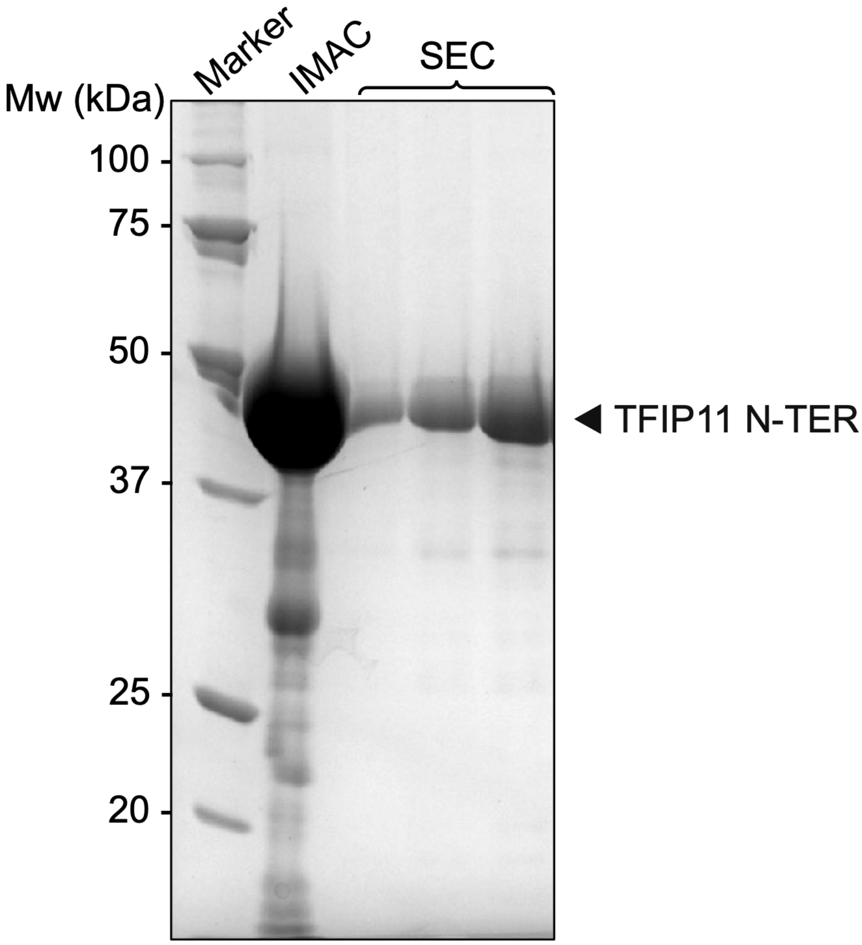


Figure S6: SDS-PAGE monitoring of His-TFIP11 N-TER purification by immobilized metal affinity chromatography (IMAC) as a first purification step (second line). The second purification step is size exclusion chromatography (SEC) (last three lines). The polyacrylamide gel is cross-linked at 12% and stained with Coomassie blue.

**TFIP11 sequence alignment**

```
sp|Q06411|SP382_YEAST      SMNSDMT--YTNDALKT-----------SSGNAPTI------SKLTKTYGIGAKLLSSMG 73
sp|Q17784|TFP11_CAEEL      SIQIDFDKRTKKAPKQNGAQVFAGMRSSANHGAADINQFGSWMRGDGNSNKIMKMMQAMG 165
sp|Q6DI35|TFP11_DANRE      -EAPPPPRAAAPKKLQTG-GSFKTSQ-RFAGGIRTGQDLGNWEKHTRGI--GQKLLQKMG 160
sp|Q9ERA6|TFP11_MOUSE      -KQEDFPKDLGPKKLKTG-GNFKPSQKGFSGGTKSFMDFGSWERHTKGI--GQKLLQKMG 162
sp|Q9UBB9|TFP11_HUMAN      -KQDDFPKDFGPRKLKTG-GNFKPSQKGFAGGTKSFMDFGSWERHTKGI--GQKLLQKMG 161
sp|Q06AK6|TFP11_PIG        -KQDEFPKDFGPKKLKTG-GNFKPSQKGFAGGTKSFMDFGSWERHTKGI--GQKLLQKMG 161
                                          :.              .           :         *::. **

sp|Q06411|SP382_YEAST      YVAGKGLGKDGSGITTPIETQSRPMHNAGLGMFSNTNSSNYHSENEDYLS---------- 123
sp|Q17784|TFP11_CAEEL      YKPGEGLGAQGQGIVEPVQAQLRKGRGA-VGAYGKESTAT---GPKFGESAADAQKRMAQ 221
sp|Q6DI35|TFP11_DANRE      YVPGKGLGKNAQGIVNPIEAKLRKGKGA-VGAYGSERTQQ---SLQDFPV-VD------- 208
sp|Q9ERA6|TFP11_MOUSE      YVPGRGLGKNAQGIINPIEAKQRKGKGA-VGAYGSERTTQ---SLQDFPV-AD------- 210
sp|Q9UBB9|TFP11_HUMAN      YVPGRGLGKNAQGIINPIEAKQRKGKGA-VGAYGSERTTQ---SMQDFPV-VD------- 209
sp|Q06AK6|TFP11_PIG        YVPGRGLGKNAQGIINPIEAKQRKGKGA-VGAYGSERTTQ---SLQDFPV-VD------- 209
                           *  *.*** :..**  *:::: *  :.* :* :..  :       :
```

Figure S7: Sequence alignment (using the multiple sequence alignment tool *Clustal Omega*) of TFIP11 analog in yeast and TFIP11 protein in different organisms. Conserved K, Y, and R residues (K155, Y162, and R166 in human TFIP11) are indicated in red boxes.

**PTM residues of TFIP11 N-TER and domain organization**

TFIP11 undergoes post-translational modification (PTM) on the residues highlighted in pink. These residues are located in different regions of the protein: K155, Y162, and R166 in the G-patch; S98 and S210 in IDR-2 and IDR-3, respectively.

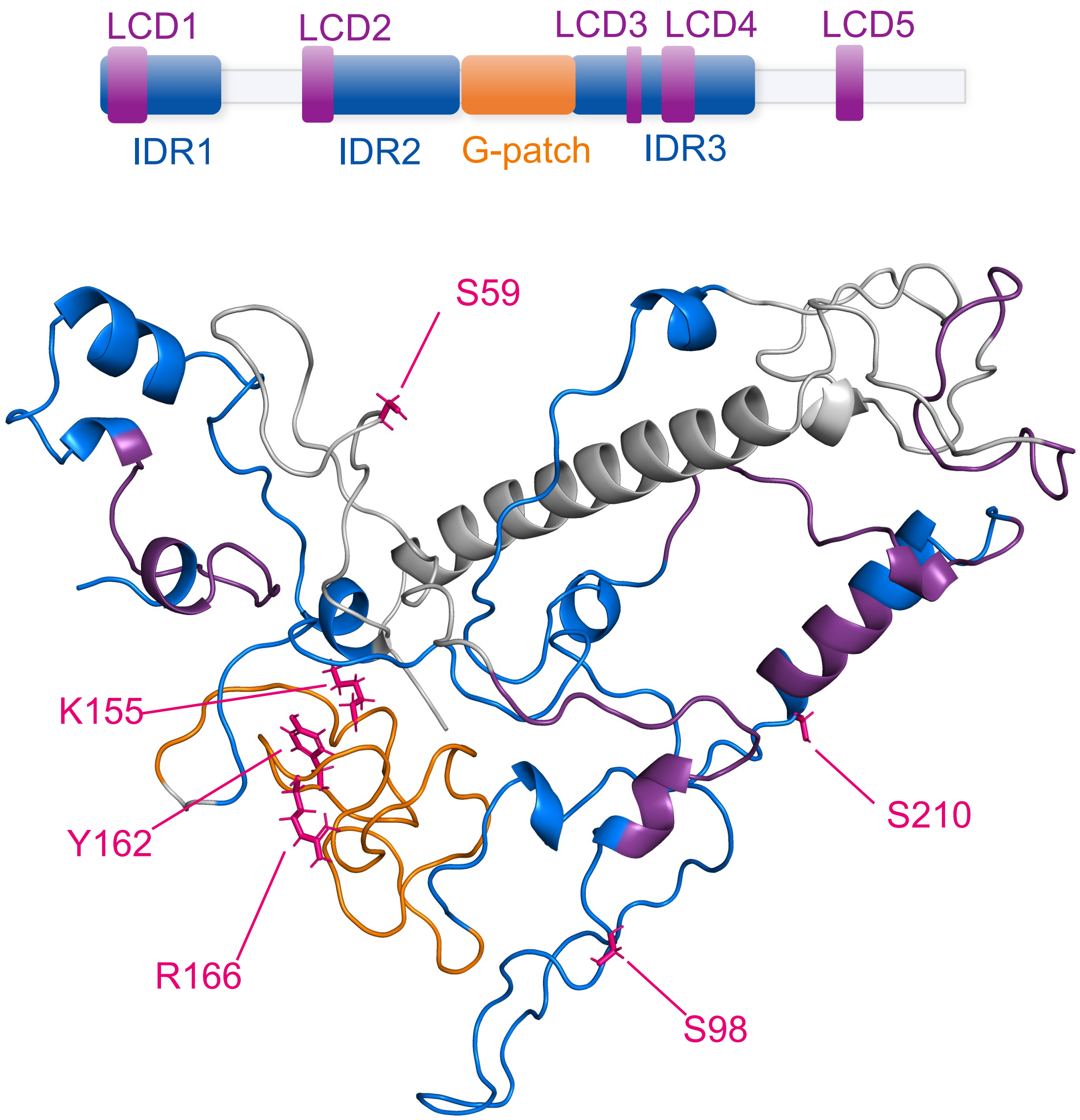


Figure S8: 3D structure of TFIP11 N-TER with the PTM residues highlighted in pink. The different domains are shown in their respective colour based on the above sequence map.

### Prediction of SLiMs

| | Positions | Séquence | Location within TFIP11 sequence | Elm description | Cell compartment |
|---|---|---|---|---|---|
| Region A | 91-95 | EEAEL | LCD2 | Fungi-specific variant of the WDR5-binding motif that binds to a cleft between blades 5 and 6 of the WD40 repeat domain of WDR5, opposite of the Win motif-binding site, to mediate assembly of histone modification complexes. | nucleus, histone methyltransferase complex |
| | 92-95 | EAEL | LCD2 | | |
| Region B | 146-152 | ERHTKGI | G-patch | Secondary preference for PKA-type AGC kinase phosphorylation | cytosol, nucleus, cAMP-dependent protein kinase complex |
| | 147-153 | RHTKGIG | G-patch | Phosphothreonine motif binding a subset of FHA domains that show a preference for a large aliphatic amino acid at the pT+3 position. | nucleus |
| | 155 -159 | KLLQK | G-patch | The USP7 CTD domain binding motif variant based on the ICP0 and DNMT1 interactions | nucleus |
| Region C | 297-303 | PLQSQQL | LCD5 | (ST)Q motif which is phosphorylated by PIKK family members. | nucleus, cytosol |
| | 297-301 | PLQSQ | LCD5 | The USP7 MATH domain binding motif variant based on the MDM2 and p53 interactions | nucleus |
| | 303-309 | LPQSGKE | LCD5 | Casein kinase 2 (CK2) phosphorylation site | protein kinase CK2 complex,nucleus,cytosol |

Figure S9: Predicted SLiMs within TFIP11 in the regions A, B, and C using the Eukaryotic Linear Motif (ELM) resource prediction tool.
